# The *Gossypium herbaceum* L. Wagad genome as a resource for understanding cotton domestication

**DOI:** 10.1101/2022.06.07.494775

**Authors:** Thiruvarangan Ramaraj, Corrinne E. Grover, Azalea C. Mendoza, Mark A. Arick, Josef J. Jareczek, Alexis G. Leach, Daniel G. Peterson, Jonathan F. Wendel, Joshua A. Udall

**Affiliations:** School of Computing, Jarvis College of Computing and Digital Media, DePaul University, Chicago, IL, 60605, USA; Ecology, Evolution, and Organismal Biology Dept., Iowa State University, Ames, IA, 50011; USDA/Agricultural Research Service, Crop Germplasm Research Unit, College Station, TX 77845; Institute for Genomics, Biocomputing & Biotechnology, Mississippi State University, United States

**Keywords:** *Gossypium herbaceum*, Wagad, genome sequence, cotton

## Abstract

*Gossypium herbaceum* is a species of cotton native to Africa and Asia that is one of the two domesticated diploids. Together with its sister-species *G. arboreum*, these A-genome taxa represent models of the extinct A-genome donor of modern polyploid cotton, which provide about 95% of cotton grown worldwide. As part of a larger effort to characterize variation and improve resources among diverse diploid and polyploid cotton genomes, we sequenced and assembled the genome of *G. herbaceum* cultivar (cv) Wagad, representing the first domesticated accession for this species. This chromosome-level genome was generated using a combination of PacBio long-read technology, HiC, and Bionano optical mapping and compared to existing genome sequences in cotton. We compare the genome of this cultivar to the existing genome of wild *G. herbaceum* subspecies *africanum* to elucidate changes in the *G. herbaceum* genome concomitant with domestication, and extend these analyses to gene expression using available RNA-seq. Our results demonstrate the utility of the *G. herbaceum* cv Wagad genome in understanding domestication in the diploid species, which could inform modern breeding programs.

## Introduction

The cotton genus (*Gossypium*) comprises the primary source of natural fiber worldwide. While the genus itself is composed of over fifty known species (Wendel and Grover 2015; Wang *et al*. 2018; Hu *et al*. 2021), only the fiber from four species is suitable for textile production. Remarkably these four species were each independently domesticated, providing a naturally replicated experiment with which we can understand the underlying genetic changes that lead to phenotypic convergence. While much research has been centered around understanding domestication in the two dominant cultivated species, i.e., *G. hirsutum* and *G. barbadense*, considerably less is known about the domestication in the “short staple cottons” (Khadi *et al*. 2010), *G. herbaceum* and *G. arboreum*.

*Gossypium herbaceum* and *G. arboreum* are diploid sister species collectively known as the “A-genome” cottons, which are native to Africa and the Arabian peninsula. Cultivated on a relatively small scale where disease and/or stress prohibit the other domesticates (Kranthi 2018), these cottons are notable both for their independent domestication (Renny-Byfield *et al*. 2016), as well as their phylogenetic relationship to the ancestral maternal parent of domesticated polyploid cottons (Wendel and Grover 2015; Huang *et al*. 2020). Accordingly, recent efforts have been made to improve resources for the A-genome cottons (Du *et al*. 2018; Huang *et al*. 2020; Grover *et al*. 2021a) in an attempt to better understand their domestication and underlying genetics. While resources are comparatively abundant for the sister species, *G. arboreum*, no wild forms have been found for that species (Vollesen 1987; Wendel *et al*. 1989a; Renny-Byfield *et al*. 2016). Native to the savannahs of Southern Africa (Vollesen 1987; Wendel *et al*. 1989a; Khadi *et al*. 2010), several accessions of wild *G. herbaceum* (subspecies *africanum*) are available from germplasm resources, as are additional accessions reflecting various levels of domestication and/or improvement.

Recently, a high-quality genome of *G. herbaceum* subspecies *africanum* has become available (Huang *et al*. 2020), providing the opportunity to evaluate the consequences of domestication in diploid cotton on a genome-wide scale. While resequencing efforts have characterized divergence of *G. herbaceum* from its sister species (*G. arboreum*) as recent (<1 million years; (Huang *et al*. 2020; Grover *et al*. 2021a)) and diversity among *G. herbaceum* accessions as low (Wendel *et al*. 1989b; Jena *et al*. 2011; Grover *et al*. 2021a), segregating polymorphisms (characterized by heterozygous, derived SNPs) number in the millions within species (1.7 - 4.1 M in each of the 21 accessions of *G. herbaceum* surveyed (Grover *et al*. 2021a)). Here we report a high-quality genome of an elite domesticated line, *G. herbaceum* cv Wagad and compare our results with the genome of wild *G. herbaceum* (subsp. *africanum*) and to the outgroup *G. longicalyx*. We present the *G. herbaceum* cv Wagad (hereafter, A1-Wagad) genome in detail, and compare this assembly to the previously released *G. herbaceum* subsp. *africanum* (hereafter, A1-africanum) assembly to provide insight into the changes accompanying domestication.

## Methods & Materials

### Plant material and sequencing methods

#### Pacbio Sequencing

Approximately 5 grams of *G. herbaceum cv Wagad* (A1-Wagad) tissue was collected from a USDA, College Station, TX greenhouse and shipped to the DNA Sequencing Center at Brigham Young University (BYU). DNA was extracted from the plant material using the Qiagen Genomic Tip kit (Qiagen, Hilden, Germany), and sequencing libraries were subsequently constructed and sequenced at the BYU DNA Sequencing Center (DNASC). DNA shearing of both libraries was done on a Megaruptor2 (∼20 kb) (Diagenonde Inc., Denville, NJ). HMW DNA was partitioned into 13 bins using the Sage Elf (Sage Science, Beverly, MA), and the top 5 bins were run on a Fragment Analyzer (Agilent Technologies, Santa Clara, CA) to select the appropriate bin size range (15-18 kb). Libraries were made using the SMRTbell Express Template Prep kit as recommended by Pacific Biosciences (PacBio; Menlo Park, CA). Three PacBio cells were sequenced from 1 CCS library on the Pacific Biosciences Sequel system.

#### Illumina Sequencing

Plant tissue of Wagad was grown at Brigham Young University Greenhouse. Young tissues were collected and DNA was extracted through the CTAB method (Kidwell and Osborn 1992). In our use of this method, tissue was lyophilized, ground in liquid nitrogen to disrupt membranes, resuspended in buffer and incubated with a lysis solution at 60° C for 30 minutes, treated with RNase, treated with chloroform to separate DNA from insoluble particles, precipitated for removal of salts, and rehydrated in TE buffer. DNA was then shipped to the National Center for Genome Resources (NCGR, Santa Fe, NM, USA) for library preparation and sequencing on the Illumina HiSeq system.

### Genome Assembly

PacBio CCS reads were assembled using Hifiasm, a fast haplotype-resolved de novo genome assembler for PacBio HiFi sequence reads (Cheng *et al*. 2021). The contigs were aligned to previously assembled genomes of *G. herbaceum* and *G. arboreum* (Huang *et al*. 2020) using minimap2 (Li 2018) and visualized by dotPlotly (https://github.com/tpoorten/dotPlotly). Multiple contigs aligning to the same contig were manually scaffolded (or concatenated) to create the final chromosomes. The final chromosomes were labeled and again aligned to the previously published A-genome using minimap2 and visualized by dotPlotly.

Hi-C libraries were constructed by Phase Genomics (Seattle, WA) from seedling tissue grown at Brigham Young University Greenhouses (different tissue source than sequenced by PacBio above). Short-read sequencing (Illumina, San Diego, CA; 150PE) of the libraries was performed by PhaseGenomics. The Hi-C data was mapped to the assembled genome sequence using bwa mem (with the -5SP option for HiC data, Li and Durbin 2009). The Hi-C interactions were used as evidence for contig proximity and in scaffolding contig sequences. Matlock (https://github.com/phasegenomics/matlock) was used in conjunction with the mapped reads to identify linkages between different genomic regions in the bam file. Juicebox (Robinson et al. 2018) was used to visualize the linkages along the pseudomolecules. All inversions were identified in alignments created by minimap2 (Li 2018).

### Repeat and gene annotation

A RepeatModeler [v2.0.1] (Flynn *et al*. 2020) generated, *Gossypium-*specific repeat database was used in conjunction with RepeatMasker [v4.1.1] (Smit *et al*. 2015a) to annotate and soft mask the repeats in the *G. herbaceum* genome. Existing RNA-seq datasets (Table S1) were mapped to the genome using hisat2 [v2.1.0] (Kim *et al*. 2015). BRAKER2 [v2.1.5] (Hoff *et al*. 2019), using a combination of GeneMark-ES [v4.61] (Borodovsky and Lomsadze 2011), the soft masked genome, and the RNA-seq alignments, was used to train Augustus [v3.3.3] (Stanke *et al*. 2006) and SNAP [v2013-02-16] (Korf 2004). The Mikado [v2.0] (Venturini *et al*. 2018) pipeline was used to predict transcripts based on the RNA-seq alignments. Briefly, the transcripts, which were predicted by Trinity [v2.11.0] (Grabherr *et al*. 2011) and mapped with GMAP [v2020-10-14] (Wu and Watanabe 2005), Cufflinks [v2.2.1] (Ghosh and Chan 2016), and Stringtie [v2.1.2] (Pertea *et al*. 2015), were combined into a non-redundant set and mapped to the Uniprot-Swissport database [v2020-05] using BLAST+ [v2.9.0] (Altschul *et al*. 1990). Additionally, Prodigal [v2.6.3] (Hyatt *et al*. 2010) was used to find ORFs in the transcripts, and Portcullis [v1.2.2] (Mapleson *et al*. 2018) was used to predict splice sites from the read alignments. Finally, Mikado used the splice site predictions, the predicted transcripts, and the protein alignments to identify gene loci and the representative transcript for each. Maker2 [v2.31.10] (Holt and Yandell 2011; Campbell *et al*. 2014) with (a) the RepeatModeler/RepeatMasker annotations; (b) the SNAP, Augustus, GeneMark, and Mikado predictions; (c) the protein evidence from previously annotated *G. hirsutum* and *G. raimondii* genomes (Chen *et al*. 2020; Paterson *et al*. 2012) along with the Uniprot Swissprot database annotations [v2020-05]; and (d) all available ESTs from NCBI for *Gossypium* (downloaded 2020-09-03, search filter: txid3633[Organism:exp] AND is_est[filter]) was used as alternative EST evidence to predict structural annotations. BUSCO [v4.1.2] (Waterhouse *et al*. 2017) was used with the eudicot [odb10] database against the predicted transcripts to find a satisfactory Annotation Edit Distance (AED) filter (Holt and Yandell 2011). An AED filter of 0.37 was used to determine the final number of genes. The filtered annotations were functionally annotated using InterProScan [v5.47-82.0] (Jones *et al*. 2014) and BLAST+ with the Uniprot Swissprot database.

### Comparison between *G. herbaceum* cv. Wagad and *G. herbaceum* subsp *africanum*

To observe syntenic conservation, whole-genome alignments were generated between wagad and africanum using Mummer (Marçais *et al*. 2018). Alignments were visualized using dotPlotly (https://github.com/tpoorten/dotPlotly) in R (version 3.6.0) (R Development Core Team and Others 2011) to identify regions with major structural variations.

Illumina DNA reads were generated and/or downloaded for A1-Wagad (PRJNA421172) and several accessions of *G. herbaceum* subspecies *africanum*, including accession ‘Mutema’ (Huang *et al*. 2020), A1-073, (Page et al. 2013), and A1-155 (Page et al. 2013). These were aligned to the outgroup reference genome *G. longicalyx* (Grover *et al*. 2020) using BWA (Li and Durbin 2009) with default arguments. Single nucleotide polymorphisms (SNPs) and small insertion-deletion polymorphisms (INDELs) were identified using the Sention (Kendig *et al*. 2019) DNAseq workflow and filtered to remove sites with >90% missing data or more than two alleles.

The resulting SNPs among A1-africanum, A1-073, A1-155, and A1-Wagad were annotated with SnpEff (Cingolani *et al*. 2012b). Since we are interested in finding SNPs that were derived under domestication (*i*.*e*., unique to A1-Wagad relative both to wild *G. herbaceum* and the outgroup reference, *G. longicalyx*) and likely visible during domestication, we filtered these SNPs using SnpSift (Cingolani *et al*. 2012a) to include only those that are homozygous variant (1/1) for A1-Wagad, homozygous reference for the three wild accessions (0/0), and predicted to be nonsynonymous (i.e., “ANN[*].EFFECT” = ‘missense_variant’, ‘exon_loss_variant’, or ‘frameshift_variant’, etc.). That is SnpSift was used to restrict the genotype for each wild accession to homozygous ancestral (e.g., GEN[africanum].GT=‘0/0’) and the A1-Wagad genotype to homozygous derived (i.e., GEN[wagad].GT=‘1/1’). Confidence in SNPs was increased via genotype level filters; *i*.*e*., Genotype Quality (GQ) less than 20 (Adelson *et al*. 2019) and Read Depth (DP) values less than 10 were removed (GEN[*].GQ > 20 & GEN[*].DP>=10).

Because fiber is one of the primary agronomic traits of interest, we screened the remaining wagad-unique nonsynonymous genes by their associated GO, Interpro, and Panther functional annotation (Grover *et al*. 2020) for those that have known functions in fiber. Any genes having homology to existing transposable elements were filtered out. To further identify and characterize changes under domestication, we used existing RNA-seq data from A1-Wagad and several *G. herbaceum* subsp. *africanum* accessions (SRA pending) to assess differential gene expression at 10 and 20 days post anthesis (DPA). RNA-seq reads for each DPA/accession were processed via Kallisto v0.46.1 (Bray *et al*. 2016) (i.e., *kallisto quant*) to identify and quantify transcripts using the A1-Wagad annotations generated here. Differential gene expression analyses were conducted in R/4.0.2 (R Core Team 2020) using DESeq2 (Love *et al*. 2014) for each DPA separately. For wild *G. herbaceum*, we used expression from multiple *G. herbaceum* subsp. *africanum* accessions as pseudoreplicates to represent expression diversity; for A1-Wagad, three replicates were used for each of the two timepoints. Genes with a Benjamini– Hochberg (Benjamini and Hochberg 1995) adjusted p-value <0.05 (as implemented in DESeq2) were considered differentially expressed.

## Data availability

The assembled genome sequence of **G. herbaceum cv. wagad** is available at NCBI under BioProject: PRJNA614591 and CottonGen (https://www.cottongen.org/). PacBio CCS sequence reads for **G. herbaceum cv. wagad** are also available at NCBI, BioProject: PRJNA421172 and (SRA pending) for RNA-Seq.

## Results and Discussion

### Genome assembly and annotation

Pacific Biosciences Sequel sequencing yielded ∼4.4 million (M) circular consensus sequence (CCS) reads with mean read length of ∼15 kbps, generating over 68.4 billion (B) bases or ≈40-fold coverage of the 1.7 gigabase (Gb) genome (Hendrix and Stewart 2005). The final assembly consisted of 13 sequences, representing the 13 chromosomes and totaling ≈1.6 Gbps (Table 1). This assembled genome size represents >94% of the estimated genome size, similar to other recent reports from *G. herbaceum* (Huang *et al*. 2020). The genome assembly was validated by Hi-C and alignment to other recent genome assemblies. The Hi-C results showed discrete colocalization along the scaffolds for cross-linked pairs of Illumina reads, with no evidence of translocations (**Figure 1**).

**Table 1.** The assembled genomes of A1-Wagad, A1-africanum, and *G. longicalyx*.

|  | *G. herbaceum cv waga* | *G. herbaceum subsp africanu* | *G. longicalyx (Outgroup)* |
| --- | --- | --- | --- |
| Contigs* | 21 | 1,782 | 97 |
| Max Contig (MB) | 141,411,576 | 9,975,897 | 49,804,510 |
| Mean Contig (MB) | 75,610,818 | 873,166 | 12,266,987 |
| Contig N50 (MB) | 122,262,091 | 1,914,893 | 29,420,374 |
| Total Contig Length | 1,587,827,183 | 1,555,981,907 | 1,189,897,770 |
| Assembly GC % | 35.1 | 35.0 | 34.2 |
| Scaffolds□ | 13 | 732 | 13 |
| Max Scaffold (MB) | 142,105,803 | 132,679,160 | 110,231,091 |
| Mean Scaffold (MB) | 122,140,578 | 2,125,802 | 91,531,230 |
| Scaffold N50 | 131,646,256 | 117,878,823 | 95,878,146 |
| Total Scaffold Length (MB) | 1,587,827,514 | 1,556,086,907 | 1,189,905,995 |
| Number of genes | 39,100 | 43,952 | 38,378 |
| *Contig metrics reports are from the raw output file of hifiasm |  |  |  |
| □ Scaffold metrics reported are after manual scaffolding |  |  |  |

**Figure 1:**
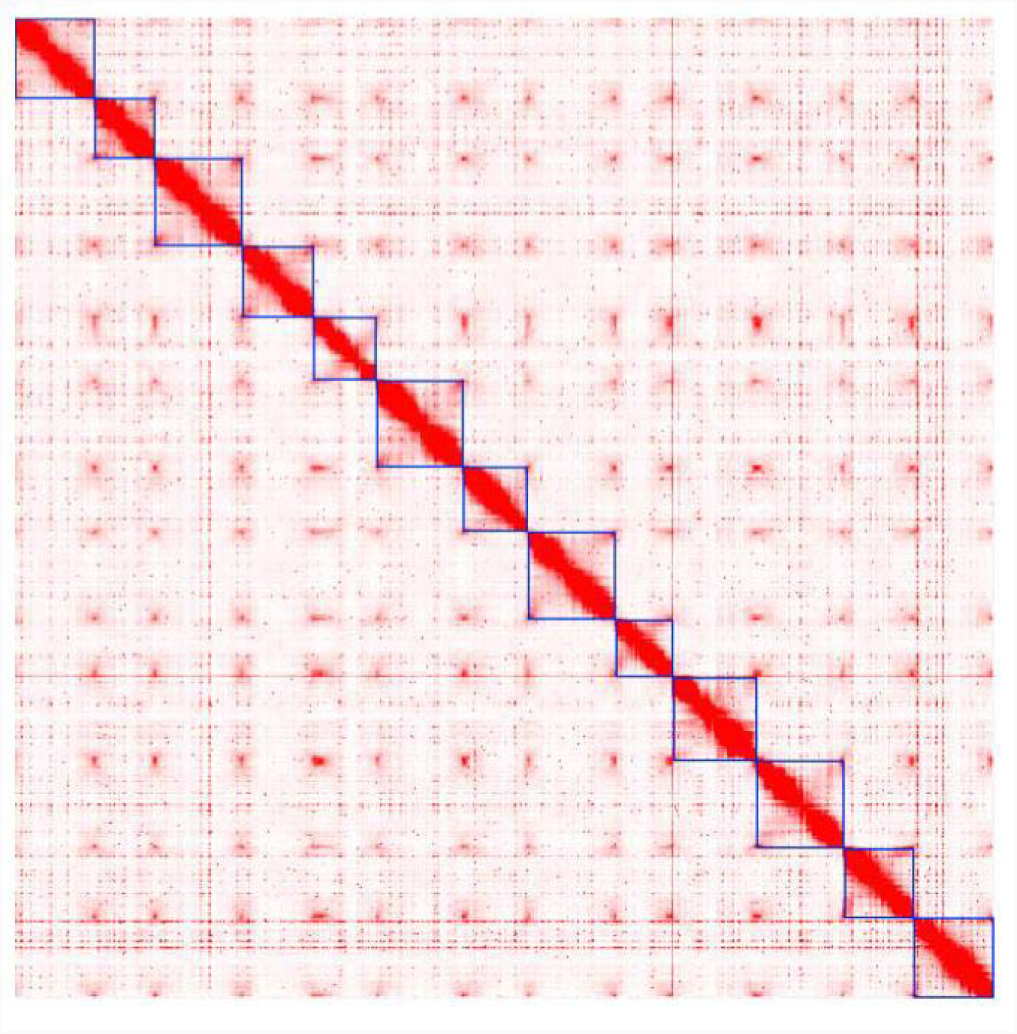
Plot of Hi-C interaction plots for the A1-Wagad genome. The x- and y-axis both depict a linear representation of the genome from top left (Chromosome 1) to bottom right (Chromosome 13). The blue boxes on the diagonal represent the chromosomes. The color intensity represents the number of HiC interactions along the chromosome and illustrates that most chromatin interactions are with other nucleotides from the same chromosome.

We used BUSCO analysis of the genome (Waterhouse *et al*. 2017) to assess the completeness of the assembly, which recovered 96.50% complete BUSCOs (from the total of 2121 BUSCO groups searched; Table 2). Most BUSCOs (88%) were both complete and single copy, with only 8.5% BUSCOs complete and duplicated. Around 3.5% of BUSCOs were either fragmented (1.20%) or missing (2.30%), indicating a general completeness of the genome. This BUSCO recovery is similar to recent publications of other diploid cotton genomes (Huang et al., 2020, Grover et al., 2020 Grover et al., 2021).

**Table 2.**
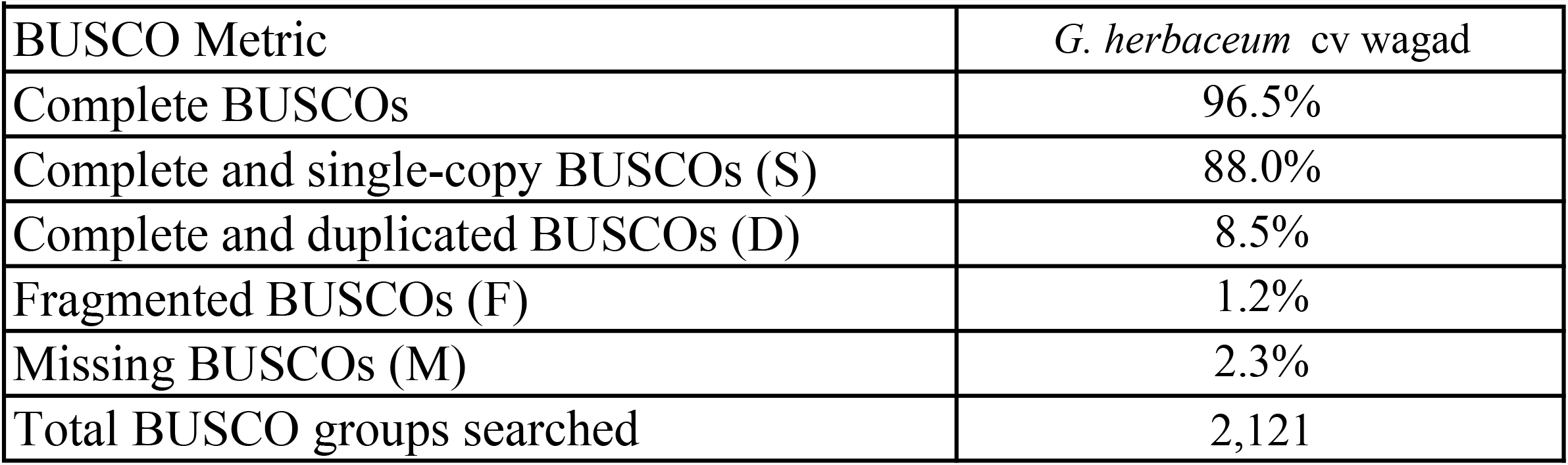
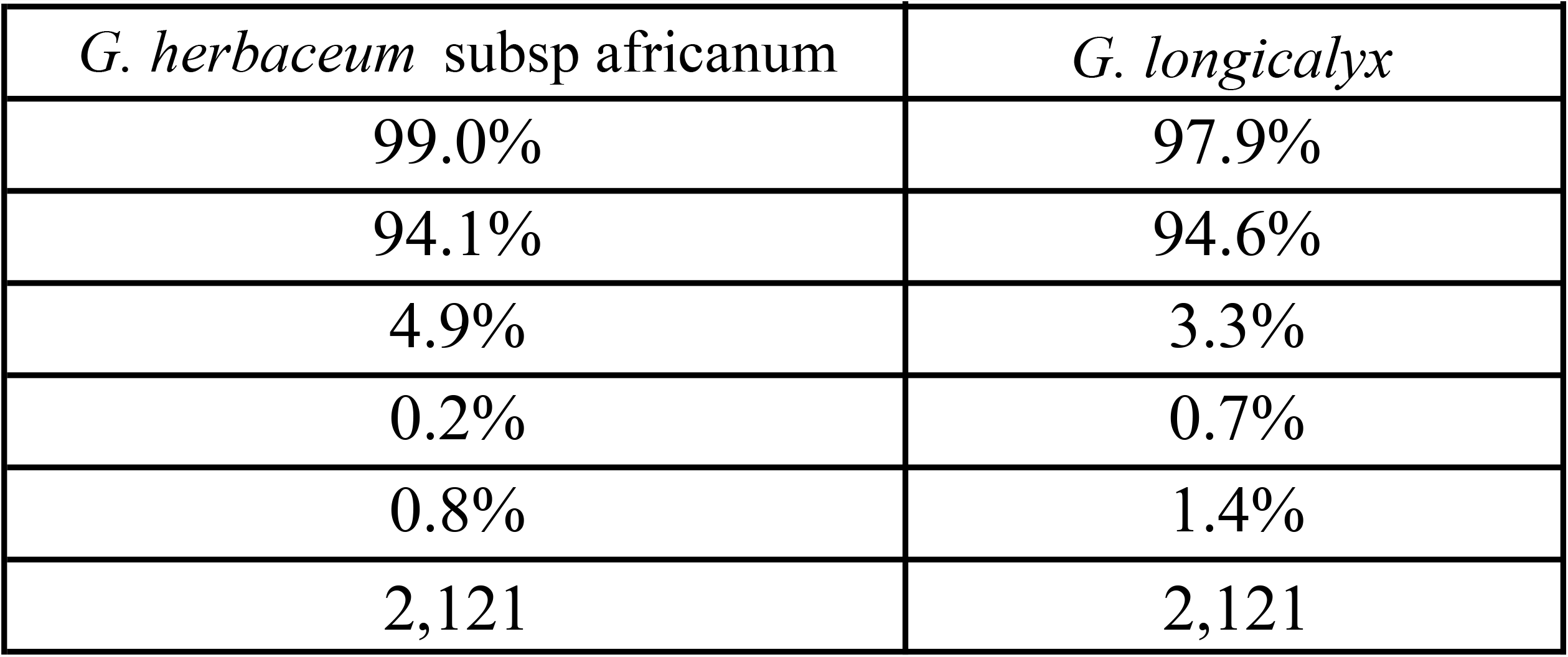
BUSCO metrics in the assembled genomes of A1-Wagad, A1-africanum, and *G. longicalyx*.

Annotation of the genome produced 39,100 unique genes, similar to the previously published cotton diploids (Paterson *et al*. 2012; Li *et al*. 2014; Du *et al*. 2018; Udall *et al*. 2019; Grover *et al*. 2020, 2021c, 2021b; Huang *et al*. 2020). The annotation of A1-Wagad contained ∼5000 fewer unique genes than the published annotation for A1-africanum, possibly due to annotation methods and/or filtering. While also fewer than previously reported for the sister taxon *G. arboreum* (40,134 to 43,278) (Li *et al*. 2014; Du *et al*. 2018; Huang *et al*. 2020), this number of annotated genes is similar to the outgroup species, *G. longicalyx* (38,378) (Grover *et al*. 2020). These discrepancies among annotations suggest there may be a future need for uniformity amongst annotation methods to further facilitate comparison among genomes.

Repeat content was predicted using a combination of RepeatMasker (Smit *et al*. 2015b), “One code to find them all” (Bailly-Bechet *et al*. 2014), and RepeatExplorer (Novák *et al*. 2013, 2020). Approximately 59.2% of the A1-Wagad genome was estimated to be repetitive, comprising 58% retroelements and 1.2% DNA transposons. As expected, more than half of the retroelements (57.3%) were annotated as LTR elements, of which 39.9% were “Gypsy/DIRS1” and 10.2% were “Ty1/Copia”; 23.3% were unclassified by the analysis [Table 3]. The repeat analysis results are very comparable to other closely related genomes, including A1-africanum, *G. arboreum, G. longicalyx*, and the A-genome of the polyploid [Table 4].

**Table 3.** Repetitive element annotations of *G. herbaceum* cv wagad

| **Family** | **Number of Elements** | **Length (Mb)** | **Percentage of Sequence** |
| --- | --- | --- | --- |
| DNA (ALL) | 21,870 | 19 | 1.22% |
| DNA/Harbinger | 4086 | 2.5 | 0.16% |
| DNA/hAT | 1,829 | 1.3 | 0.08% |
| LTR (ALL) | 1,011,698 | 921 | 58.01% |
| LTR | 1,001,971 | 909 | 57.31% |
| LTR/Copia | 281,785 | 161 | 10.18% |
| LTR/Gypsy | 519,278 | 633 | 39.87% |
| EVERYTHING_TE |  | 1,310 | 82.51% |

**Table 4.**
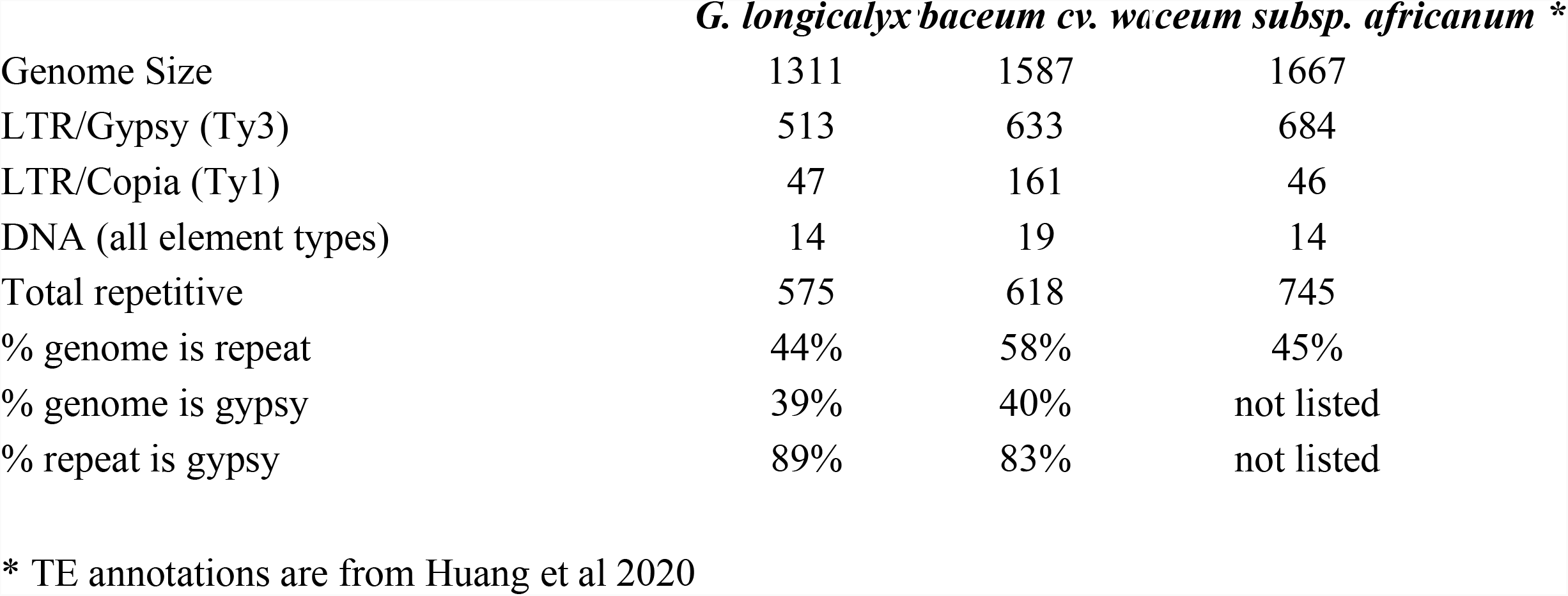
Repeat element content in A1-Wagad versus A1-africanum and *G. longicalyx*

### Comparison between cultivated A1-wagad and wild A1-africanum

Domestication is an important evolutionary process whereby human-mediated phenotypic changes result in myriad (often anonymous) corresponding molecular changes. Understanding the molecular basis for improved phenotypes is a fundamental goal of many modern breeding programs, which may use a combination of traditional and cutting edge breeding techniques to accelerate crop improvement. Comparison between wild and domesticated germplasm can improve our understanding of the molecular basis underlying the phenotypic superiority of modern crops, as well as facilitate our understanding of how wild germplasm can be used to improve modern cultivars (e.g., by conferring disease resistance).

Whole genome alignment of the A1-Wagad sequence to the recent assembly of A1-africanum revealed high co-linearity with a few exceptions, including major inversions (≈6 M) on A1-01, A1-02, and A1-06 and two inversions on A1-13 (≈43 M; **Figure 2 and Figures Supp 1-6**). When these two genomes were compared to the genome of their sister species *G. arboreum* (Huang *et al*. 2020; Wang *et al*. 2021), we found many shared inversions between *G. herbaceum* genome sequences (A1-africanum and A1-Wagad) and the *G. arboreum* genome sequence. The shared inversions and high degree of co-linearity suggested that the independent *G. herbaceum* genome sequences were correctly assembled. In one case, however, we found a unique inversion in the A1-Wagad sequence on A1-06 (102,159,887-110,192,639), possibly indicating a structural change acquired during or after domestication (**Supp Figures 1-6**).

**Figure 2.**
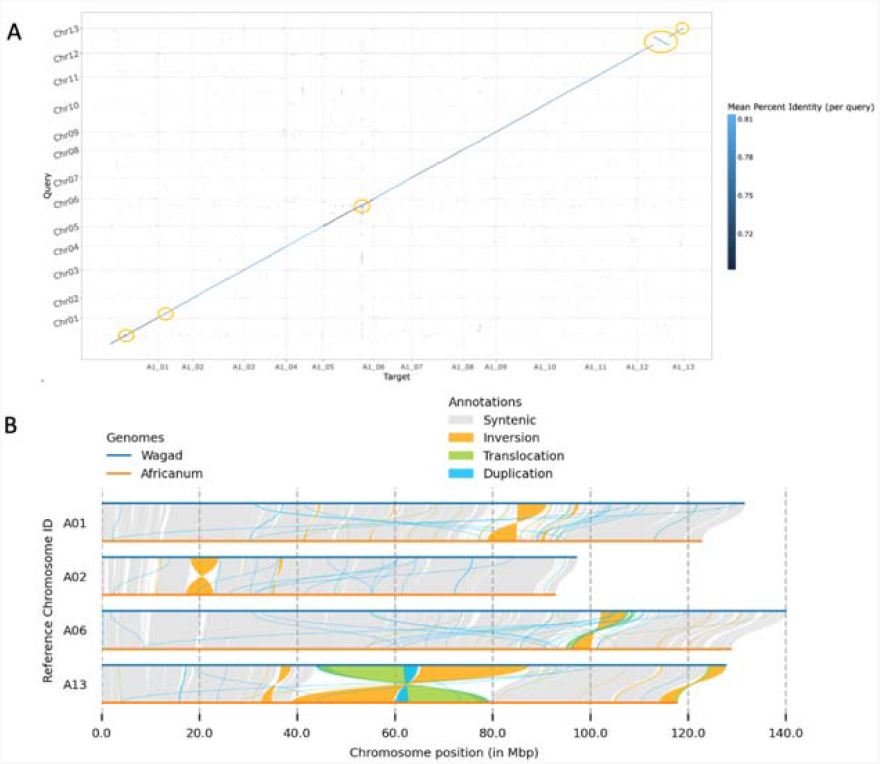
A) Genome alignment of the 13 chromosomes in the G. herbaceum var africanum (y-axis) and the G. herbaceum cv Wagad (x-axis) genomes. The shade of blue indicates the degree of sequence match. The orange circles indicate major and minor inversions between the two genome assemblies on chromosomes A01, A02, A06, and A13. B) Alignment of the chromosomes containing inversions (A01, A02, A06, and A13) between the two A1-genomes. A13 had 3 inversions. The major inversion on A13 was so large that a portion was colored as a translocation, instead of an inversion by plotsr (Goel and Schneeberger 2022).

At the nucleotide level, we identified 220,191 fixed SNPs between A1-Wagad and the three A1-africanum accessions. As expected, 84% of the SNPs were intergenic, and only 17% were within gene regions (including UTR). Because amino acid changes may be visible to selection (*i*.*e*., domestication), we focused on derived nonsynonymous mutations in A1-Wagad; that is, A1-Wagad was homozygous for a derived SNP where all three A1-africanum accessions shared the ancestral state with *G. longicalyx*. This produced a set of 4,719 nonsynonymous SNPs that occurred in 2,544 genes. Notably, 26 of these had >10 missense mutations (accounting for 433 SNPs), likely indicating a major disruption to the gene, such as a frameshift mutation and/or pseudogenization. Many of these genes also had other variant annotations (e.g., UTR, start/stop codon gain/loss, splice site modifications), while an additional 779 genes had at least one of these other potentially functionally relevant annotations without also having a missense mutation. Overall, the number of genes with only single, missense mutation was 524 or ∼25% of the total, indicating that fixed differences between wild and domesticated *G. herbaceum*, while affecting <10% of the genes, tend to produce multiple modifications. A full list of genes, their inferred variant effects, their *G. herbaceum* homologs, and their functional annotations is listed in Supplementary Table 3.

We also leveraged existing RNA-seq data to infer transcriptomic changes between wild and domesticated *G. herbaceum* fiber at two key timepoints, *i*.*e*., 10 and 20 DPA, that are representative of primary and secondary wall synthesis, respectively. Here, RNA-seq was mapped against the gene annotations of the newly developed A1-Wagad sequence to infer differential expression (Table DE; Supplementary Table 1). For 10 DPA, 6034 genes were differentially expressed (3175 down, 2859 up), including 990 genes that were also on the list of genes with annotated SNP variants (Supplementary Table 2). Functional annotation of these genes include genes with possible relevance to fiber, such as three upregulated tubulin-related proteins (A1_06G097900, A1_05G253000, A1_10G076200) and several upregulated kinesin-like proteins (Supplementary Table 2). These two examples are important for fiber shape, as tubulins comprise the shape conferring microtubules while kinesins move materials along them (Graham and Haigler 2021; Preuss et al. 2004). Likewise, at 20 DPA 6400 genes exhibited differential expression (3078 down, 3322 up), including 1040 genes exhibiting potentially consequential SNP variants. Among these are several downregulated NAC domain-containing proteins, which regulate secondary cell wall deposition (Zhong et al. 2006; Zhang et al. 2018), and four probable WRKY transcription factors, most of which are also downregulated, with the exception of A1_09G185500, which is upregulated 2.7-fold in A1-Wagad. There were 1979 genes differentially expressed at both timepoints, most of which exhibited either upregulation (736, or 37%) or downregulation (647, 33%) at both timepoints. For those genes that exhibited differences in relative expression level across timepoints, 445 were downregulated at 10 DPA but upregulated at 20 DPA, and only 151 genes exhibited the inverse (i.e., upregulated in A1-Wagad at 10 DPA, but downregulated at 20 DPA). A similar pattern was observed for the 355 genes that also contained an annotated variant (Supplementary Table 2). These results provide an overview of differential expression in wild versus domesticated diploid cotton fiber using the newly generated A1-Wagad genome.

## Conclusion

Cotton is an important fiber crop that has been independently domesticated multiple times. While the cultivated tetraploid species have extensive genome resources, cultivated diploid cotton species have received far less attention. Here we report a genome sequence for a cultivated form of *Gossypium herbaceum* (cv Wagad) that complements the existing genome assemblies and diversity studies (Huang *et al*. 2020). Our data provide a foundation for understanding the transition from wild to domesticated *G. herbaceum*, thereby providing an additional perspective on the commonalities of domestication in multiple cotton species.

## Acknowledgements

We thank the National Science Foundation (NSF #1339412) and USDA-ARS (58-6066-0-066, Genomics of Malvaceae) for their financial support. This work was made possible by the USDA Supercomputer Atlas funded through the SCINet initiative. We thank the Iowa State University ResearchIT unit for computational resources and support.

## References

Adelson, R. P., A. E. Renton, W. Li, N. Barzilai, G. Atzmon et al., 2019 Empirical design of a variant quality control pipeline for whole genome sequencing data using replicate discordance. Sci. Rep. 9: 16156.

Altschul, S. F., W. Gish, W. Miller, E. W. Myers, and D. J. Lipman, 1990 Basic local alignment search tool. J. Mol. Biol. 215: 403–410.

Bailly-Bechet, M., A. Haudry, and E. Lerat, 2014 “One code to find them all”: a perl tool to conveniently parse RepeatMasker output files. Mob. DNA 5: 13.

Benjamini, Y., and Y. Hochberg, 1995 Controlling the False Discovery Rate: A practical and powerful approach to multiple testing. J. R. Stat. Soc. Series B Stat. Methodol. 57: 289–300.

Borodovsky, M., and A. Lomsadze, 2011 Eukaryotic gene prediction using GeneMark.hmm-E and GeneMark-ES. Curr. Protoc. Bioinformatics Chapter 4: Unit 4.6.1–10.

Bray, N. L., H. Pimentel, P. Melsted, and L. Pachter, 2016 Erratum: Near-optimal probabilistic RNA-seq quantification. Nat. Biotechnol. 34: 888.

Campbell, M. S., C. Holt, B. Moore, and M. Yandell, 2014 Genome Annotation and Curation Using MAKER and MAKER-P. Curr. Protoc. Bioinformatics 48: 4.11.1–39.

Cheng, H., G. T. Concepcion, X. Feng, H. Zhang, and H. Li, 2021 Haplotype-resolved de novo assembly using phased assembly graphs with hifiasm. Nat. Methods 18: 170–175.

Cingolani, P., V. M. Patel, M. Coon, T. Nguyen, S. J. Land et al., 2012a Using Drosophila melanogaster as a Model for Genotoxic Chemical Mutational Studies with a New Program, SnpSift. Front. Genet. 3: 35.

Cingolani, P., A. Platts, L. L. Wang, M. Coon, T. Nguyen et al., 2012b A program for annotating and predicting the effects of single nucleotide polymorphisms, SnpEff: SNPs in the genome of Drosophila melanogaster strain w1118; iso-2; iso-3. Fly 6: 80–92.

Du, X., G. Huang, S. He, Z. Yang, G. Sun et al., 2018 Resequencing of 243 diploid cotton accessions based on an updated A genome identifies the genetic basis of key agronomic traits. Nat. Genet. 50: 796–802.

Flynn, J. M., R. Hubley, C. Goubert, J. Rosen, A. G. Clark et al., 2020 RepeatModeler2 for automated genomic discovery of transposable element families. Proc. Natl. Acad. Sci. U. S. A.

Ghosh, S., and C.-K. K. Chan, 2016 Analysis of RNA-Seq Data Using TopHat and Cufflinks. Methods Mol. Biol. 1374: 339–361.

Grabherr, M. G., B. J. Haas, M. Yassour, J. Z. Levin, D. A. Thompson et al., 2011 Full-length transcriptome assembly from RNA-Seq data without a reference genome. Nat. Biotechnol. 29: 644–652.

Grover, C. E., M. A. Arick, A. Thrash, J. Sharbrough, G. Hu et al., 2021a Dual domestication, diversity, and differential introgression in Old World cotton diploids. bioRxiv 2021.10.20.465142.

Grover, C. E., M. Pan, D. Yuan, M. A. Arick, G. Hu et al., 2020 The Gossypium longicalyx Genome as a Resource for Cotton Breeding and Evolution. G3.

Grover, C. E., D. Yuan, M. A. Arick, E. R. Miller, G. Hu et al., 2021b The Gossypium anomalum genome as a resource for cotton improvement and evolutionary analysis of hybrid incompatibility. G3.

Grover, C. E., D. Yuan, M. A. Arick, E. R. Miller, G. Hu et al., 2021c The Gossypium stocksii genome as a novel resource for cotton improvement. G3.

Hendrix, B., and J. M. Stewart, 2005 Estimation of the nuclear DNA content of gossypium species. Ann. Bot. 95: 789–797.

Hoff, K. J., A. Lomsadze, M. Borodovsky, and M. Stanke, 2019 Whole-Genome Annotation with BRAKER. Methods Mol. Biol. 1962: 65–95.

Holt, C., and M. Yandell, 2011 MAKER2: an annotation pipeline and genome-database management tool for second-generation genome projects. BMC Bioinformatics 12: 491.

Huang, G., Z. Wu, R. G. Percy, M. Bai, Y. Li et al., 2020 Genome sequence of Gossypium herbaceum and genome updates of Gossypium arboreum and Gossypium hirsutum provide insights into cotton A-genome evolution. Nat. Genet. 52: 516–524.

Hu, G., C. E. Grover, D. Yuan, Y. Dong, E. Miller et al., 2021 Evolution and Diversity of the Cotton Genome, pp. 25–78 in Cotton Precision Breeding, edited by M.-U.- Rahman, Y. Zafar, and T. Zhang. Springer International Publishing, Cham.

Hyatt, D., G.-L. Chen, P. F. Locascio, M. L. Land, F. W. Larimer et al., 2010 Prodigal: prokaryotic gene recognition and translation initiation site identification. BMC Bioinformatics 11: 119.

Jena, S. N., A. Srivastava, U. M. Singh, S. Roy, N. Banerjee et al., 2011 Analysis of genetic diversity, population structure and linkage disequilibrium in elite cotton (Gossypium L.) germplasm in India. Crop Pasture Sci. 62: 859–875.

Jones, P., D. Binns, H.-Y. Chang, M. Fraser, W. Li et al., 2014 InterProScan 5: genome-scale protein function classification. Bioinformatics 30: 1236–1240.

Kendig, K. I., S. Baheti, M. A. Bockol, T. M. Drucker, S. N. Hart et al., 2019 Sentieon DNASeq Variant Calling Workflow Demonstrates Strong Computational Performance and Accuracy. Front. Genet. 10: 736.

Khadi, B. M., V. Santhy, and M. S. Yadav, 2010 Cotton: An Introduction, pp. 1–14 in Cotton: Biotechnological Advances, Springer Berlin Heidelberg, Berlin, Heidelberg.

Kidwell, K. K., and T. C. Osborn, 1992 Simple plant DNA isolation procedures, pp. 1–13 in Plant Genomes: Methods for Genetic and Physical Mapping, edited by J. S. Beckmann and T.C. Osborn. Springer Netherlands, Dordrecht.

Kim, D., B. Langmead, and S. L. Salzberg, 2015 HISAT: a fast spliced aligner with low memory requirements. Nat. Methods 12: 357–360.

Korf, I., 2004 Gene finding in novel genomes. BMC Bioinformatics 5: 59.

Kranthi, K. R., 2018 Cotton production practices: snippets from global data 2017. The ICAC Recorder XXXVI: 4–14.

Li, H., and R. Durbin, 2009 Fast and accurate short read alignment with Burrows-Wheeler transform. Bioinformatics 25: 1754–1760.

Li, F., G. Fan, K. Wang, F. Sun, Y. Yuan et al., 2014 Genome sequence of the cultivated cotton Gossypium arboreum. Nat. Genet. 46: 567–572.

Love, M. I., W. Huber, and S. Anders, 2014 Moderated estimation of fold change and dispersion for RNA-seq data with DESeq2. Genome Biol. 15: 550.

Mapleson, D., L. Venturini, G. Kaithakottil, and D. Swarbreck, 2018 Efficient and accurate detection of splice junctions from RNA-seq with Portcullis. Gigascience 7.:

Marçais, G., A. L. Delcher, A. M. Phillippy, R. Coston, S. L. Salzberg et al., 2018 MUMmer4: A fast and versatile genome alignment system. PLoS Comput. Biol. 14: e1005944.

Novák, P., P. Neumann, and J. Macas, 2020 Global analysis of repetitive DNA from unassembled sequence reads using RepeatExplorer2. Nat. Protoc. 15: 3745–3776.

Novák, P., P. Neumann, J. Pech, J. Steinhaisl, and J. Macas, 2013 RepeatExplorer: a Galaxy-based web server for genome-wide characterization of eukaryotic repetitive elements from next-generation sequence reads. Bioinformatics 29: 792–793.

Paterson, A. H., J. F. Wendel, H. Gundlach, H. Guo, J. Jenkins et al., 2012 Repeated polyploidization of Gossypium genomes and the evolution of spinnable cotton fibres. Nature 492: 423–427.

Pertea, M., G. M. Pertea, C. M. Antonescu, T.-C. Chang, J. T. Mendell et al., 2015 StringTie enables improved reconstruction of a transcriptome from RNA-seq reads. Nat. Biotechnol. 33: 290–295.

R Core Team, 2020 R: A language and environment for statistical computing. R Foundation for Statistical Computing., Vienna, Austria.

R Development Core Team, R., and Others, 2011 R: A language and environment for statistical computing.

Renny-Byfield, S., J. T. Page, J. A. Udall, W. S. Sanders, D. G. Peterson et al., 2016 Independent Domestication of Two Old World Cotton Species. Genome Biol. Evol. 8: 1940–1947.

Smit, A. F. A., R. Hubley, and P. Green, 2015a RepeatMasker Open-4.0. 2013--2015.

Smit, A. F. A., R. Hubley, and P. Green, 2015b RepeatModeler Open-1.0. 2008--2015. Seattle, USA: Institute for Systems Biology. Available from: http://www.repeatmasker.org, Last Accessed May 1: 2018.

Stanke, M., O. Keller, I. Gunduz, A. Hayes, S. Waack et al., 2006 AUGUSTUS: ab initio prediction of alternative transcripts. Nucleic Acids Res. 34: W435–9.

Udall, J. A., E. Long, C. Hanson, D. Yuan, T. Ramaraj et al., 2019 De Novo Genome Sequence Assemblies of Gossypium raimondii and Gossypium turneri. G3 9: 3079–3085.

Venturini, L., S. Caim, G. G. Kaithakottil, D. L. Mapleson, and D. Swarbreck, 2018 Leveraging multiple transcriptome assembly methods for improved gene structure annotation. Gigascience 7.:

Vollesen, K., 1987 The Native Species of Gossypium (Malvaceae) in Africa, Arabia and Pakistan. Kew Bull. 42: 337–349.

Wang, M., J. Li, P. Wang, F. Liu, Z. Liu et al., 2021 Comparative Genome Analyses Highlight Transposon-Mediated Genome Expansion and the Evolutionary Architecture of 3D Genomic Folding in Cotton. Molecular Biology and Evolution.

Wang, K., J. F. Wendel, and J. Hua, 2018 Designations for individual genomes and chromosomes in Gossypium. Journal of Cotton Research 1: 3.

Waterhouse, R. M., M. Seppey, F. A. Simão, M. Manni, P. Ioannidis et al., 2017 BUSCO applications from quality assessments to gene prediction and phylogenomics. Mol. Biol. Evol.

Wendel, J. F., and C. E. Grover, 2015 Taxonomy and Evolution of the Cotton Genus, Gossypium, pp. p25–44 in Cotton, edited by D. D. Fang and R.G. Percy. Agronomy Monographs, American Society of Agronomy, Inc., Crop Science Society of America, Inc., and Soil Science Society of America, Inc., Madison, WI, USA.

Wendel, J. F., P. D. Olson, and J. M. Stewart, 1989a GENETIC DIVERSITY, INTROGRESSION, AND INDEPENDENT DOMESTICATION OF OLD WORLD CULTIVATED COTTONS. Am. J. Bot. 76: 1795–1806.

Wendel, J. F., P. D. Olson, and J. M. Stewart, 1989b Genetic diversity, introgression, and independent domestication of Old World cultivated cottons. Am. J. Bot. 76: 1795–1806.

Wu, T. D., and C. K. Watanabe, 2005 GMAP: a genomic mapping and alignment program for mRNA and EST sequences. Bioinformatics 21: 1859–1875.

