## Supplemental Figures 1 - 6 for "The *Gossypium herbaceum* L. Wagad genome as a resource for understanding cotton domestication"

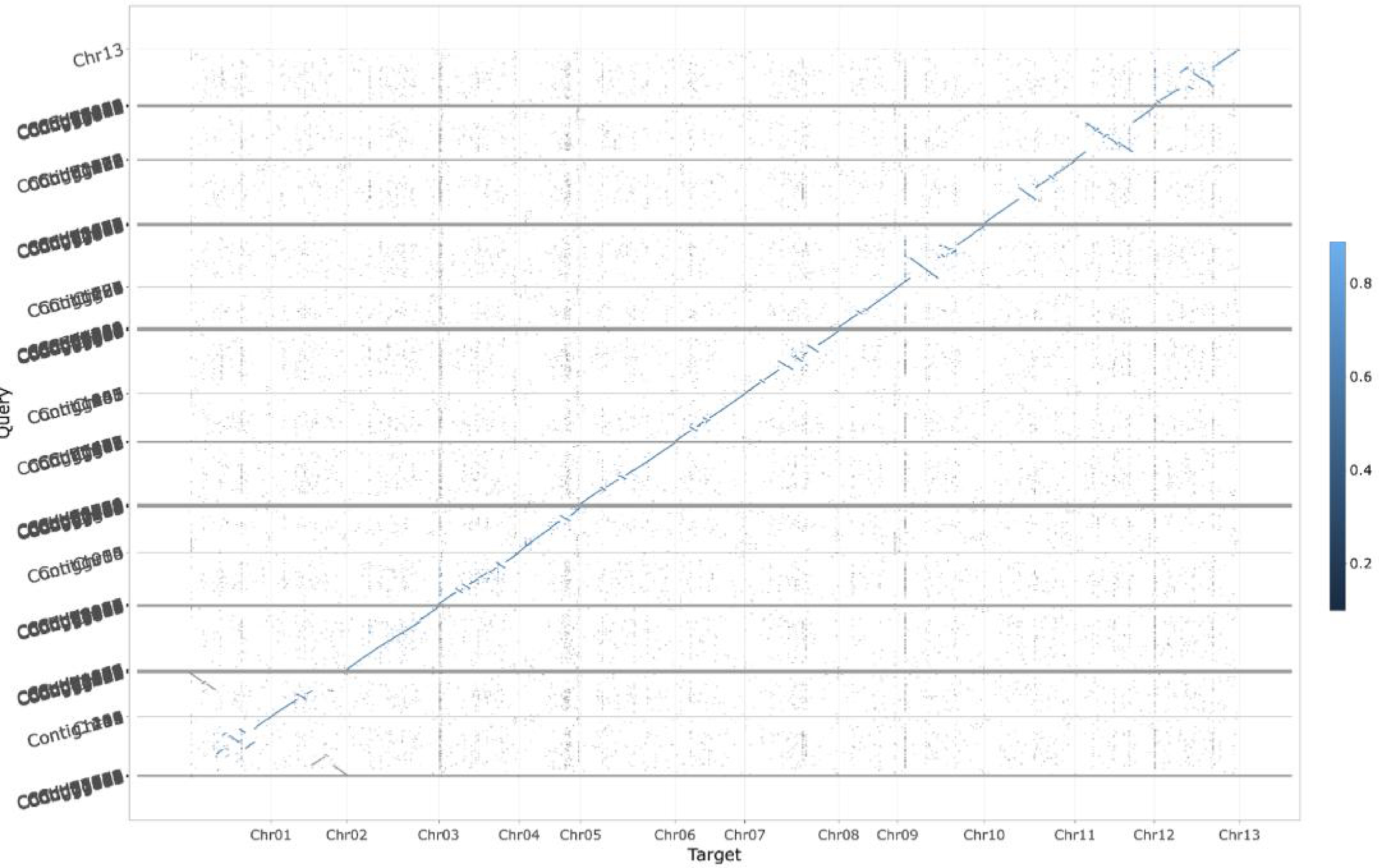


Supplementary Figure 1. An alignment between the G. arboreum cv Shi Xi Ya 1 (x-axis, Query, Huang et al. 2020) and the G. herbaceum var. africanum (y-axis, Target) genome sequences. Many inversions are observed through-out the genome particularly an inversion on A1, A2,  and the two inversions on A13 (Supp Figures 3-5). These are unique to the G. herbaceum var. africanum genome

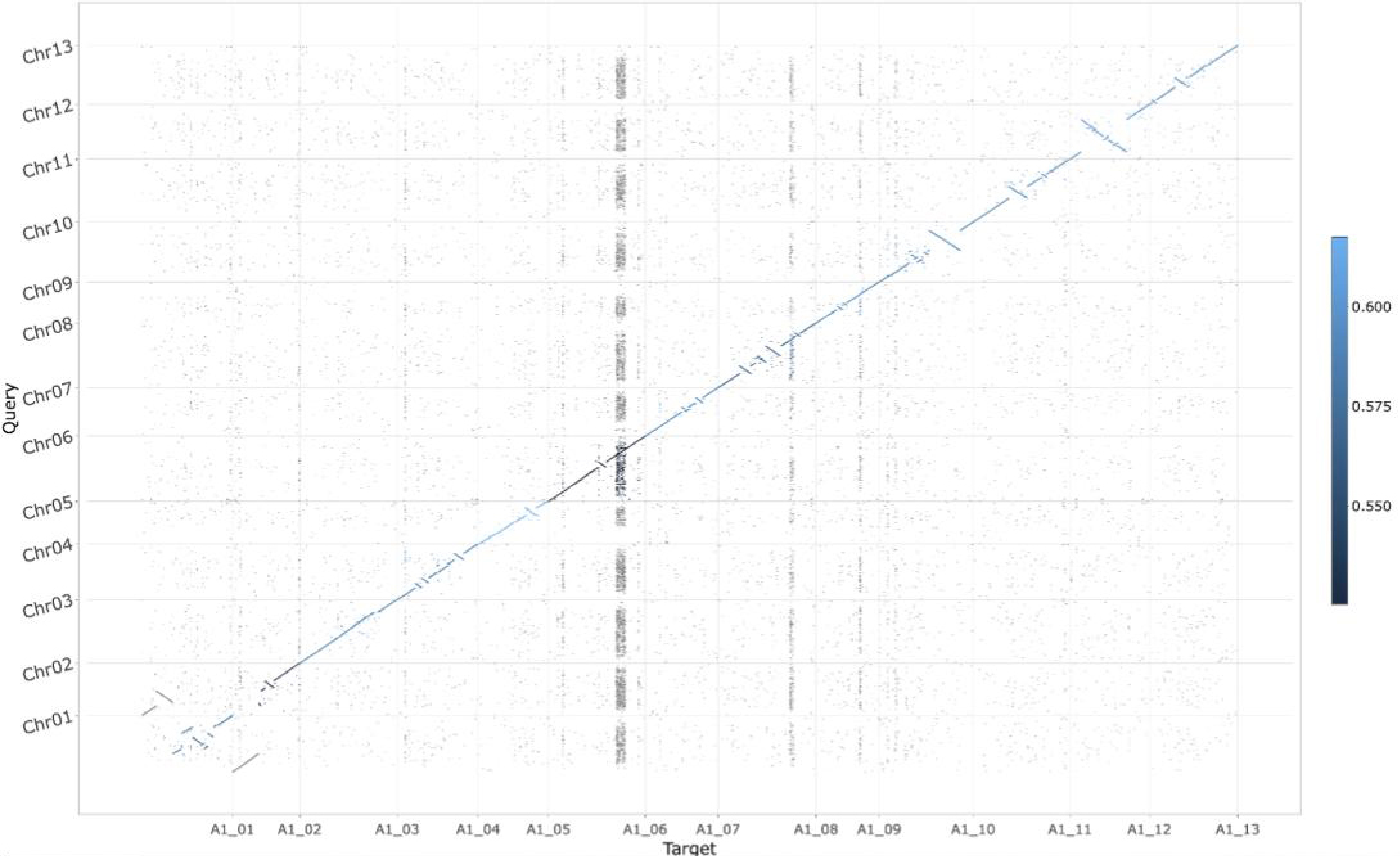


Supplemental Figure 2. An alignment between the the G. herbaceum cv Wagad (x-axis, Target) and the G. arboreum cv Shi Xi Ya 1 (y-axis, Query, Huang et al. 2020) genome sequences. The inversion on A06 is distinct in G. herbaceum cv Wagad

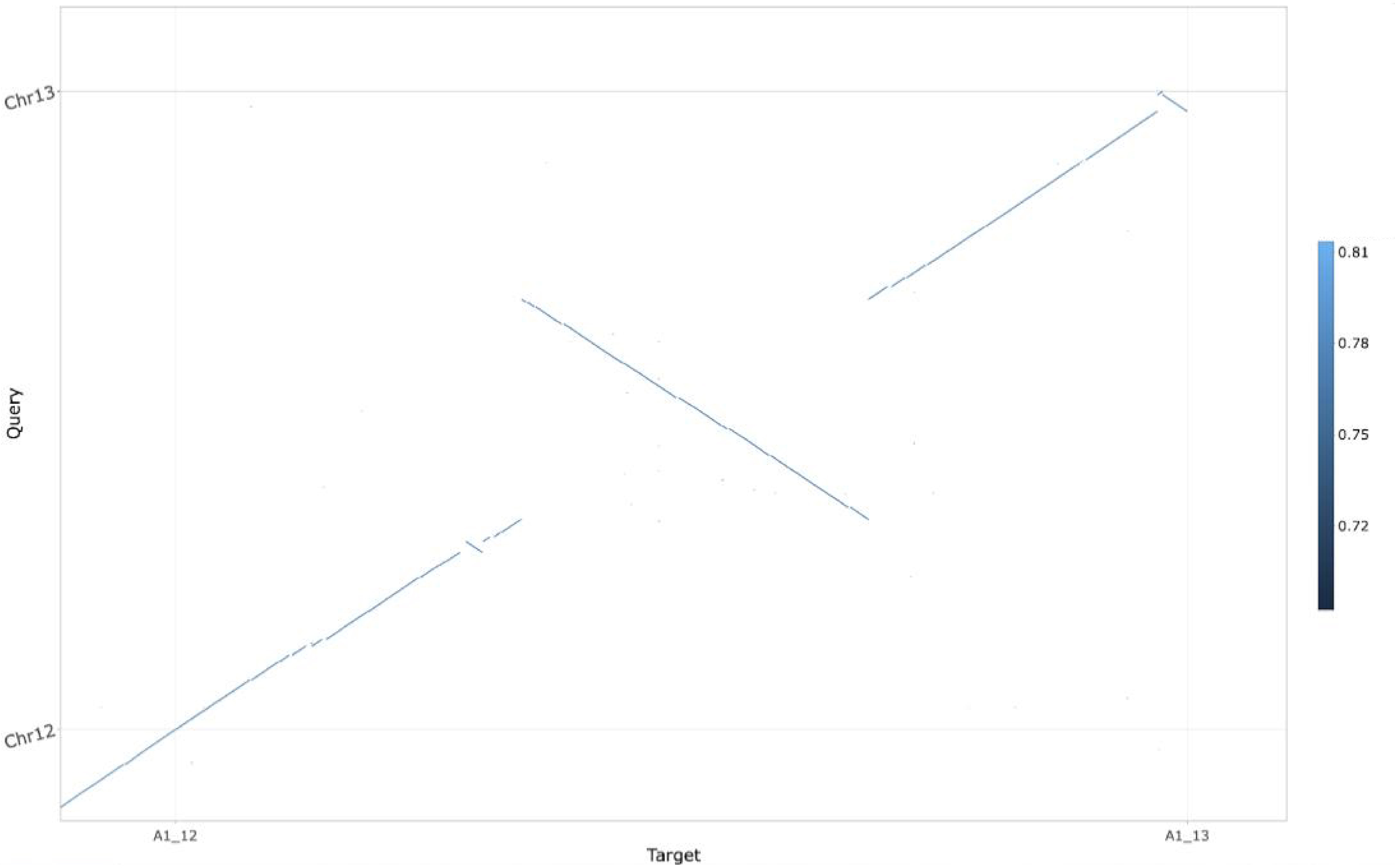


Supplemental Figure 3. An alignment of Chr12 between the G. herbaceum cv Wagad (x-axis, Target) and G. herbaceum var. africanum (y-axis, Query) genome sequences. Two large inversions are found on A12 - one in the middle and one one the end.

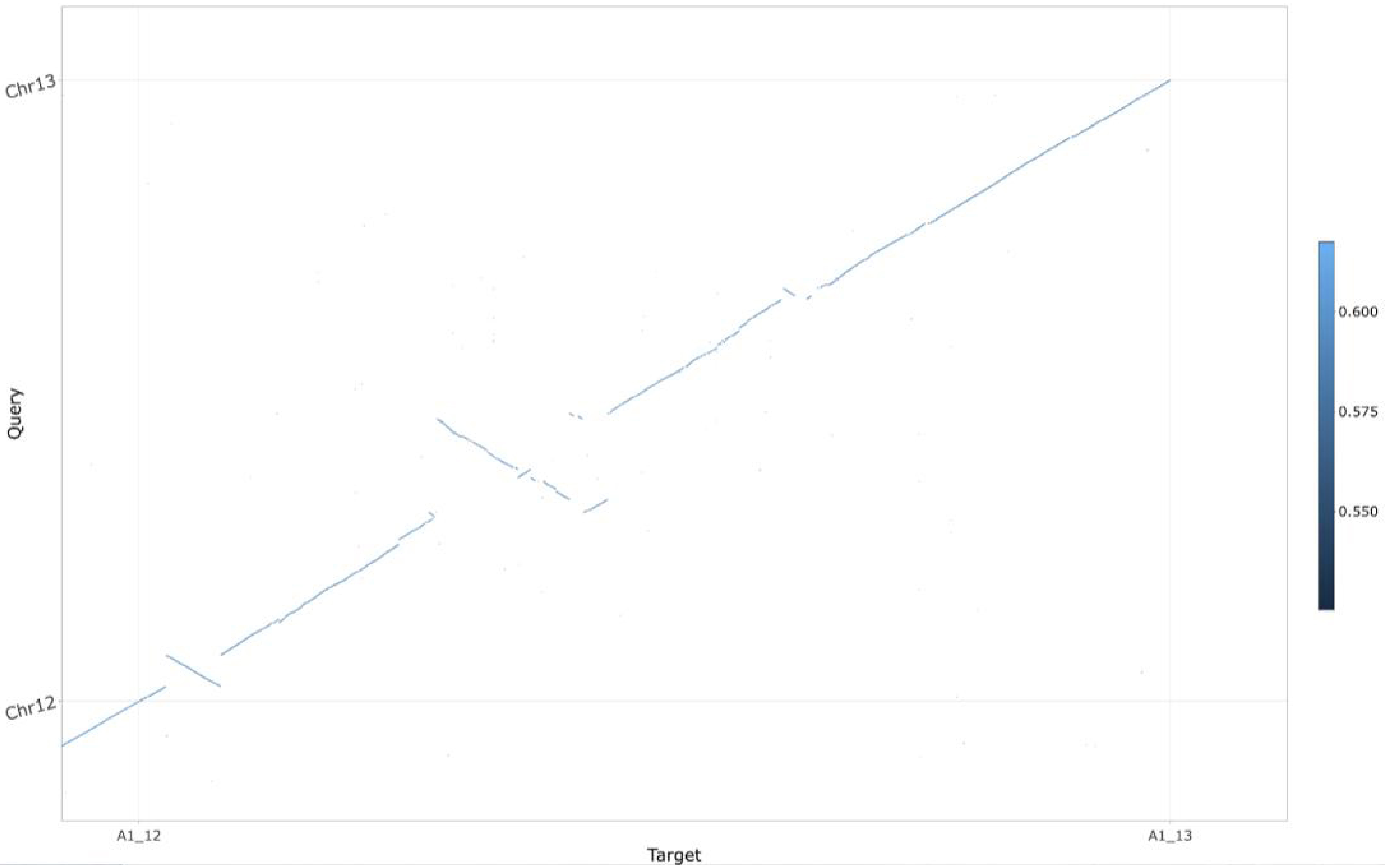


Supplemental Figure 4. An alignment between the G. herbaceum Wagad (x-axis, Target) and the G. arboreum cv Shi Xi Ya 1 (y-axis, Query, Huang et al. 2020) genome sequences. The pictured inversion is different size than the the inversion between the G. herbaceum cv Wagad and G. herbaceum var. africanum 'Mutema' (Supp Fig 4). The two inversions seen here (large central inversion and a modest distal inversion) are found in both A1-genome vs. A2-genome comparisons (Wagad vs. Shi Xi Ya 1 (Supp Fig 5); Mutema vs. Shi Xi Ya 1 (Supp Fig 6)) suggesting that these inversions are found in both Wagad and Mutema and occurred prior other genome rearrangements between the different A1 genomes (Wagad and Mutema).

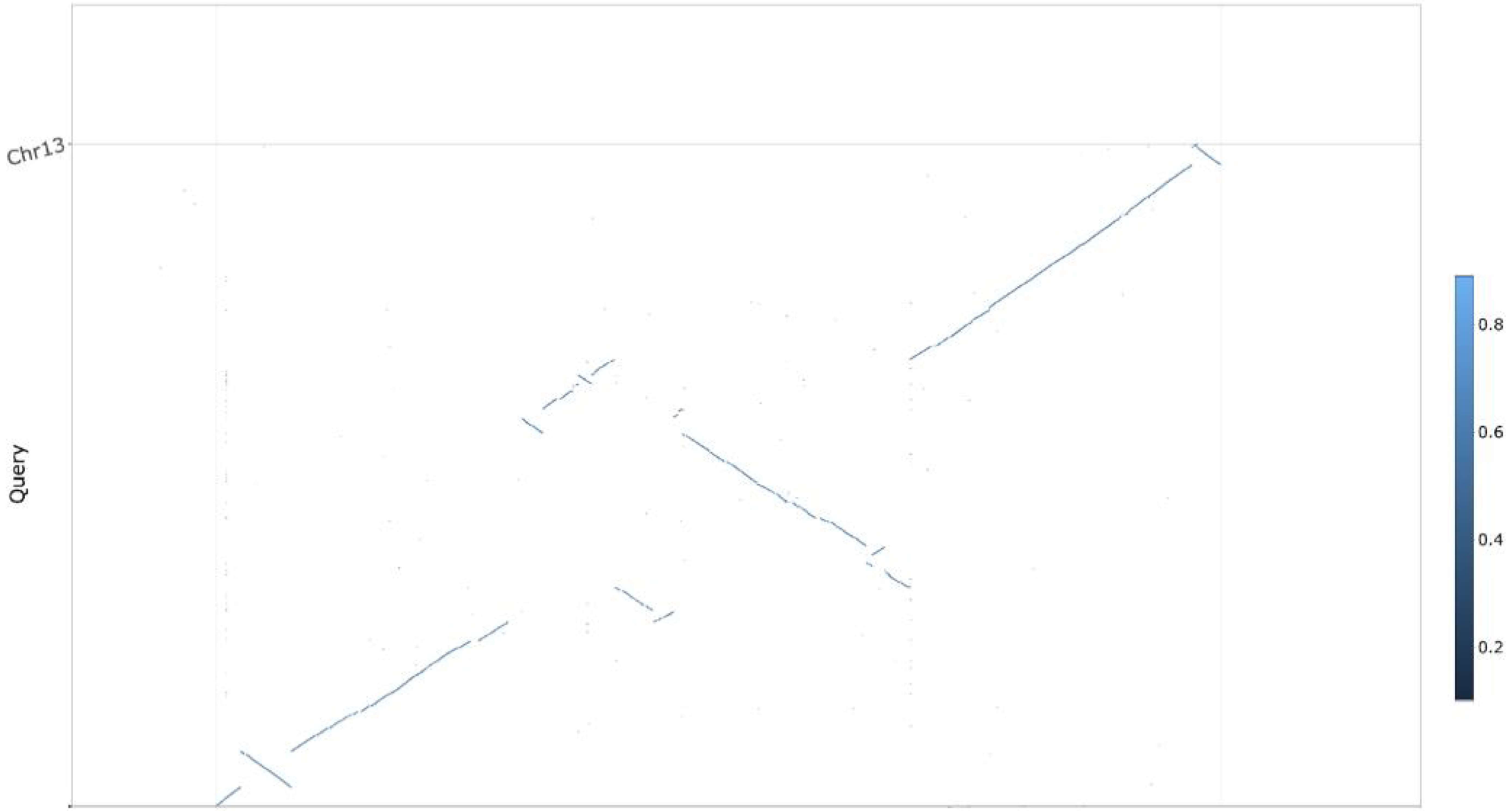


Supplemental Figure 5. An alignment between the G. arboreum cv Shi Xi Ya 1 (x-axis, Query, Huang et al. 2020) and the G. herbaceum var. africanum 'Mutema' (y-axis, Target) genome sequences. The large inversion between A1 and A2 genomes (Supp Fig 4) happened after the inversions pictured in Supp Fig 5. This resulted in an inversion (Supp Fig 4) of an inversion (Supp Fig 5) in the central region to create the composite pattern seed in the alignment of these two genomes.

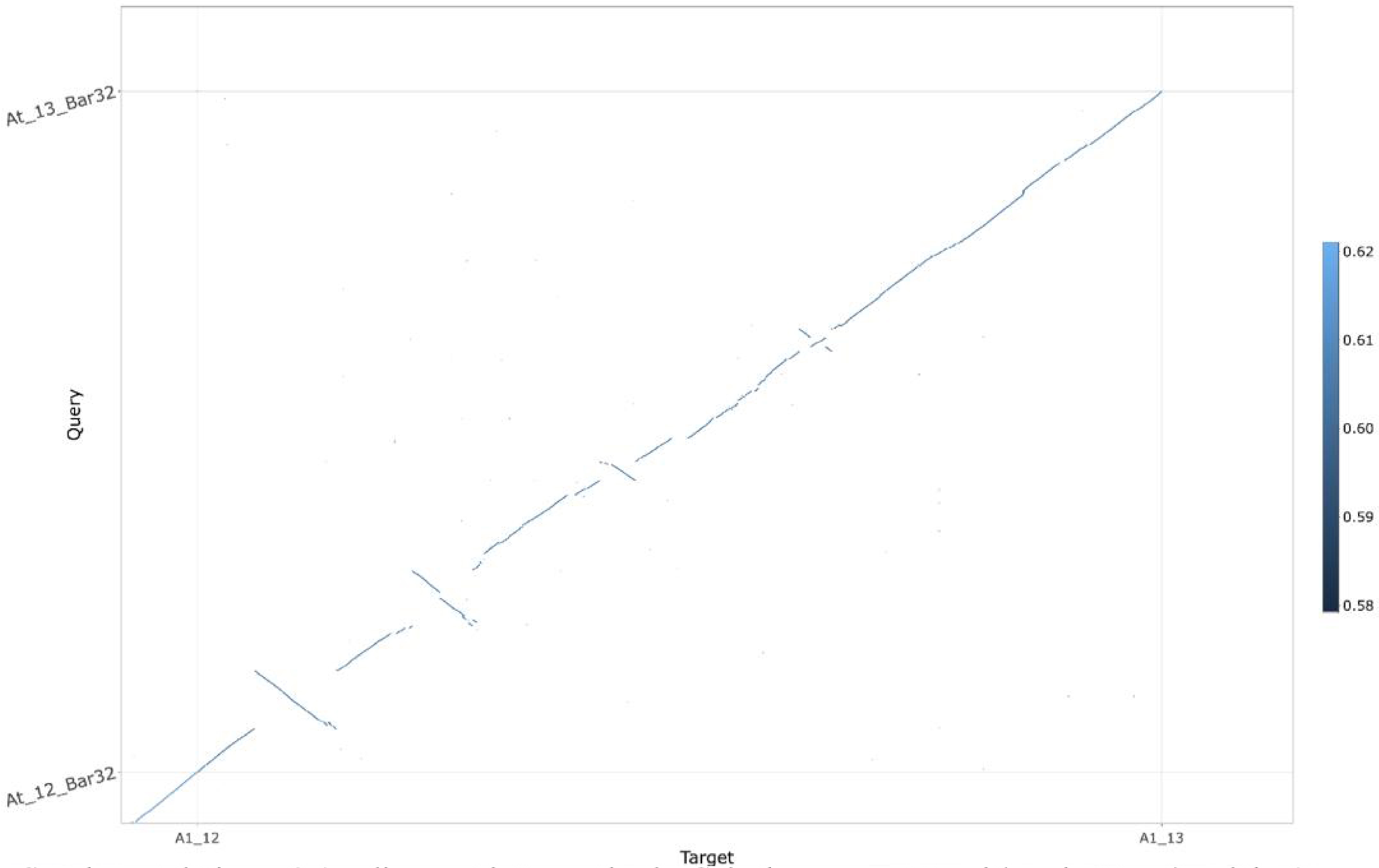


Supplemental Figure 6. An alignment between the the G. herbaceum cv Wagad (x-axis, Target) and the A-genome found in tetraploid cotton G. hirsutum (Perkin et al. 2021).
